# Tracing the expansion of p53 retrogenes in elephant species: A foundation for functional insights

**DOI:** 10.64898/2026.02.25.707932

**Authors:** Konstantinos Karakostis, Elena Campoy, Marta Puig, Robin Fåhraeus, Fritz Vollrath, Mario Cáceres

**Affiliations:** Biochemistry and Molecular Biology Department, University of Valencia, Spain; Research Programme on Biomedical Informatics (GRIB), Hospital del Mar Research Institute, Barcelona, Spain; Institut de Biotecnologia i de Biomedicina, Universitat Autònoma de Barcelona, Bellaterra (Barcelona), Spain; Departament de Genètica i de Microbiologia, Universitat Autònoma de Barcelona, Bellaterra (Barcelona), Spain; Inserm UMRS1131, Institut de Génétique Moléculaire, Université Paris 7, Hôpital St. Louis, Paris, France; Research Centre for Applied Molecular Oncology (RECAMO), Masaryk Memorial Cancer Institute, Brno, Czech Republic; Department of Medical Biosciences, Umeå University, Umeå, Sweden; Department of Biology, University of Oxford, Oxford, UK; Save the Elephants Marula Manor, Marula Lane, Karen P.O. Box 54667, Nairobi, Kenya; ICREA, Barcelona, Spain

**Keywords:** tumour suppressor, cancer, genomic evolution, p53, Peto’s Paradox, afrotheria

## Abstract

Elephants have evolved multiple *TP53* copies through a retrotransposition event followed by successive duplications. Some of these *TP53* retrogenes (RTGs) are expressed and hypothesized to have functional roles in cellular regulation. However, comparative genomic studies on *TP53* evolution and function are limited due to scarce genomic data for elephants and other afrotherians. Most existing research relies on scaffold assemblies of *Loxodonta africana* (LoxAfr3 and LoxAfr4), with some focus on *Elephas maximus* chromosomal assembly. In this *in silico* study, we analyzed three elephant genomes to validate *TP53* RTGs, assess their copy variation, and trace their evolution. For the first time we describe 29 *TP53* RTGs in *E. maximus* versus 18-19 in *L. africana*. These copies show sequence variation, especially in the duplicated regions and their flanking repetitive elements. Chromosomal mapping in *E. maximus* revealed that two major classes of *TP53* RTGs are consistently arranged in pairs on chromosome 27, which harbours 27 of the 29 identified copies. The observed distribution strongly supports an evolutionary model in which large-scale genomic segments, each encompassing at least two retrogenes of different groups, were duplicated early in the elephant lineage, driving the extensive amplification of *TP53* RTGs, as suggested also by the patterns of the flanking repetitive elements. The *TP53* RTGs expansion was followed by a unique inversion on chromosome 27 that separates the duplication clusters. Thus, this study enhances our understanding of the elephant’s multi-p53 system, linked to cancer resistance, body size, and Peto’s paradox, and supports ongoing research into functional aspects.

**Significance Statement:** This study uncovers the complex genomic architecture of elephant *TP53* RTGs, which have diversified into two phylogenetically distinct types present in both African and Asian elephants. While both species share these 2 types, they differ substantially in copy number (18 in African versus 28 in Asian elephants), and in sequence variation, highlighting lineage-specific evolutionary trajectories. Detailed sequence analyses of the chromosomal organization of these copies in the *Elephas maximus* high-quality genome assembly, particularly the cluster on chromosome 27, indicates that they arose from stepwise duplications of extended genomic segments, which likely facilitated both the expansion and regulatory diversification of the *TP53* RTGs repertoire. Comparative analysis with other mammals reveals an elephant-specific inversion that reorganized the expanded *TP53* RTGs copies, creating a new genomic configuration that could have influenced the regulation and expression of some of the *p53* RTGs. Therefore, this work advances our understanding of how evolutionary pressures shaped the landscape of *TP53* RTGs and their flanking genetics elements in elephants. By identifying accurately the retrogene copy numbers, sequences and putative functional domains, it establishes a critical foundation for future studies investigating their functional roles in DNA damage and genome stability.

## Introduction

The ATM–MDM2–p53 axis is the central pathway regulating p53 activity [1–6]. At the heart of this regulation is the p53 transactivation domain, whose conserved BOX-I motif mediates binding to the E3 ubiquitin ligase MDM2. This interaction has co-evolved at both the protein and mRNA levels [7, 8], highlighting its essential role in p53 activation. Even though the MDM2/p53 regulatory interaction is currently the most promising antitumor p53 target [9, 10], interventions targeting this interaction have not reached therapies mainly due to its? dynamic structural complexity [5]. Comparative studies with divergent model systems expressing modified homologous p53 and MDM2 encoding genes can provide potentially groundbreaking insights at both the structural and signaling mechanisms that lead to p53 activation. Interestingly, elephants carry dozens of *TP53* retrogene (RTG) copies of variable sequences and yet poorly described functions [11–13]. Some of these variants display different MDM2 binding affinities and activation capacities, pointing towards a lineage-specific diversification of this critical pathway [14, 15]

Throughout evolution there are very few examples of positive selection leading to increased copy number of transcription factors linked to cancer, increased body size or extended lifespan [16]. Specifically, recent studies on *L. africana* have revealed 18 *TP53* RTGs partial p53-coding sequences [11, 12, 17]. These are proposed to derive from a single retroposition event, also present in manatee and hyrax, followed by repeated rounds of segmental duplications in the elephant [18]. The flanking regions of the *TP53* duplicated sequences contain different transposable elements (TEs) [12], including several retrotransposons. Such retroposition processes typically generate non-functional pseudogenes. However, their amplification and long-term maintenance could reflect a combination of neutral retention and functional co-option, as some of these retrogenes have already been shown to play roles in apoptosis in mitochondria [13]. Therefore, while most *TP53* RTG copies may be non-functional, a subset could have been retained under selective pressure due to lineage-specific contributions to genome stability and stress responses [19].

Several independent studies have aimed to understand the evolution of the *TP53* copies and their potential functions. In particular, their coding potential was evaluated on the basis of scaffold-level genomic assemblies of *L. africana* (LoxAfr3 and LoxAfr4 [11, 12, 20]. An independent study on LoxAfr3 investigated the extent of the conservation of the p53 coding sequence and the potential transcription of each RTG. Those results indicated that the RTGs derive from duplications of a truncated ancestral elephant RTG (157 conserved aa out of the initial 390 aa in the original p53), caused by a frameshift mutation and argued that 14 of the 18 copies are shown to presently be truncated to ≤88 aa, without evidence of positive or negative selection [18]. However, several studies provide experimental evidence of *p53* RTGs functional implications. *TP53* RTG12 was shown to enhance p53 signalling and DNA-damage response, while co-immunoprecipitation experiments on transiently transfected *HEK-293* cells demonstrated a lack of interaction between RTG12 and MDM2, indicating that it escapes negative regulation [12]. These results support the “guardian model”, whereby the products of certain elephant *TP53* RTGs interact with the canonical p53 forming polymers with specific functions [12]. In another example, monomeric *TP53* RTG9 was shown to uptake independent functions and induce transcription-independent apoptosis at the mitochondria [13]. Functional aspects were further investigated by focusing on variations of the BOX-I motif, which is well conserved throughout evolution and constitutes the primary binding epitope that regulates the negative regulation of p53 by MDM2 [14]. The nineteen p53(BOX-I) motifs of LoxAfr3 *TP53* RTGs were categorised into seven types (A to F) by testing their binding to MDM2, both by *in silico* docking models and *in vitro* with synthetic BOX-I peptides. Results showed that the BOX-I peptides exhibit a spectrum of binding affinities to MDM2 with potential functional roles and an integrative model was proposed accounting for a range of activities of the truncated copies in response to various stresses [14].

However, the current methodologies employ partially assembled genomic sequences with low coverage across the identified *TP53* copies (ie, the 7X-coverage LoxAfr3 scaffold genome assembly) and heterologous cell systems (ie, transfected human cell lines), resulting in incomplete conclusions [11]. Overcoming such limitations is necessary to understand the potential functional aspects of the multiple *TP53* copies in the elephant system and their contribution to p53 regulation. In this study we aimed to resolve discrepancies in *TP53* RTG number and sequences by leveraging the chromosome-level and higher-quality assembly of the Asian elephant genome, and investigate the evolution of the duplication events. We specifically asked: (i) how many *TP53* RTG can be confidently identified in the chromosome-level assembly of the Asian elephant, and how do these compare with previously reported numbers in African and Asian elephants, based on lower-quality scaffold assemblies? (ii) Does their sequence variation reflect assembly accuracy or population-level polymorphism or lineage-specific divergence? (iii) how their flanking sequences and phylogeny inform the evolutionary trajectory of *TP53* RTG expansion? By providing this precise genomic framework, we establish a foundation for functional studies, such as performing CRISPR/Cas9 KO, identifying stress-responsive promoters or exploring specific interactions of MDM2 with those similar but not identical BOX-I motifs of each *TP53* RTG [14].

## Materials and Methods

### Alignments of TP53-copies and flanking regions in three genomes

**Supplementary Table 1** summarizes the main characteristics of the genomes used in this study and their accession numbers. The canonical African elephant (*L. africana*) *TP53* coding sequence (CDS) XP_010594888.1 was aligned with BLAST v2.4.0 [21] against the same species genome assemblies LoxAfr3 (GCF_000001905.1) and LloxAfr4 [20] (available at ftp://broadin-stitute.org/pub/assemblies/mammals/elephant/loxAfr4/). The 18 identified sequences in each genome were highly similar to the previous GenBank entries *TP53* RTG 1 to 18 (KF715855.1 to KF715872.1), and they were named accordingly **(Supplementary Table 2)**. These sequences were aligned and plotted in pairs **(Supplementary Table 3)**. Similarly, the same canonical *TP53* CDS was also aligned with BLAST against the Asian elephant (*E. maximus*) chromosome level genome assembly (GCF_024166365.1, mEleMax1) **(Supplementary Table 2,4)**. In this genome, 29 *TP53* RTGs were identified, which for simplicity were labeled with successive numbers (1 to 27) based on their position on Chr. 27 with 28 and 29 being in two other chromosomes, and their sequences were also aligned and plotted in pairs **(Supplementary Table 4)**. Phylogenetic trees were constructed using IQ-TREE (version 2.2.0) [22] following the alignment of target sequences using Clustal Omega (version 1.2.4) [23]. Tree construction was performed with the command: “*iqtree2 -s $alignmentFile -m MFP -B 1000 -alrt 1000”*, which involved selecting the best-fitting nucleotide substitution model, performing 1000 bootstrap replicated, and conducting 1000 approximate likelihood ratio test (aLRT) replicates for branch support. We constructed two different phylogenetic trees. The first shows the relationship between the canonical *TP53* and *TP53* RTGs sequences of *E. maximus, E. telfairi, Hyracoidea sp.* and *Trichechus sp*. **(Figure 1A)** while the second also includes the corresponding sequences for *L. africana* (**Supplementary Figure 1, Supplementary Table 4B**). Next, flanking sequences extended bilaterally by 50 kb were extracted from the genome using bedtools getfasta v2.31.0 [24]. Dot-Plot analysis was performed to identify the extended alignments and to reveal groups of *TP53* RTG-bearing sequences that show higher identity **(Figure 1B)**. Circus Plots were generated using the R package “Biocircos” v.0.3.4 [25] to illustrate the alignments and identity among the duplicated extended *TP53* sequences **(Figure 1C).** Codon aware alignments were also generated using MACSE (version 2.07). The align Sequences option was used to translate nucleotide sequences into amino acids, perform alignment at the protein level, and subsequently reconstruct codon-aware nucleotide alignments. The resulting sequences were visualised using Clustal Omega (version 1.2.4) and phylogenetic trees were inferred. We observed no differences in the groupings of the sequences compared to the non-codon aware alignments (Data not shown).

**Figure 1.**
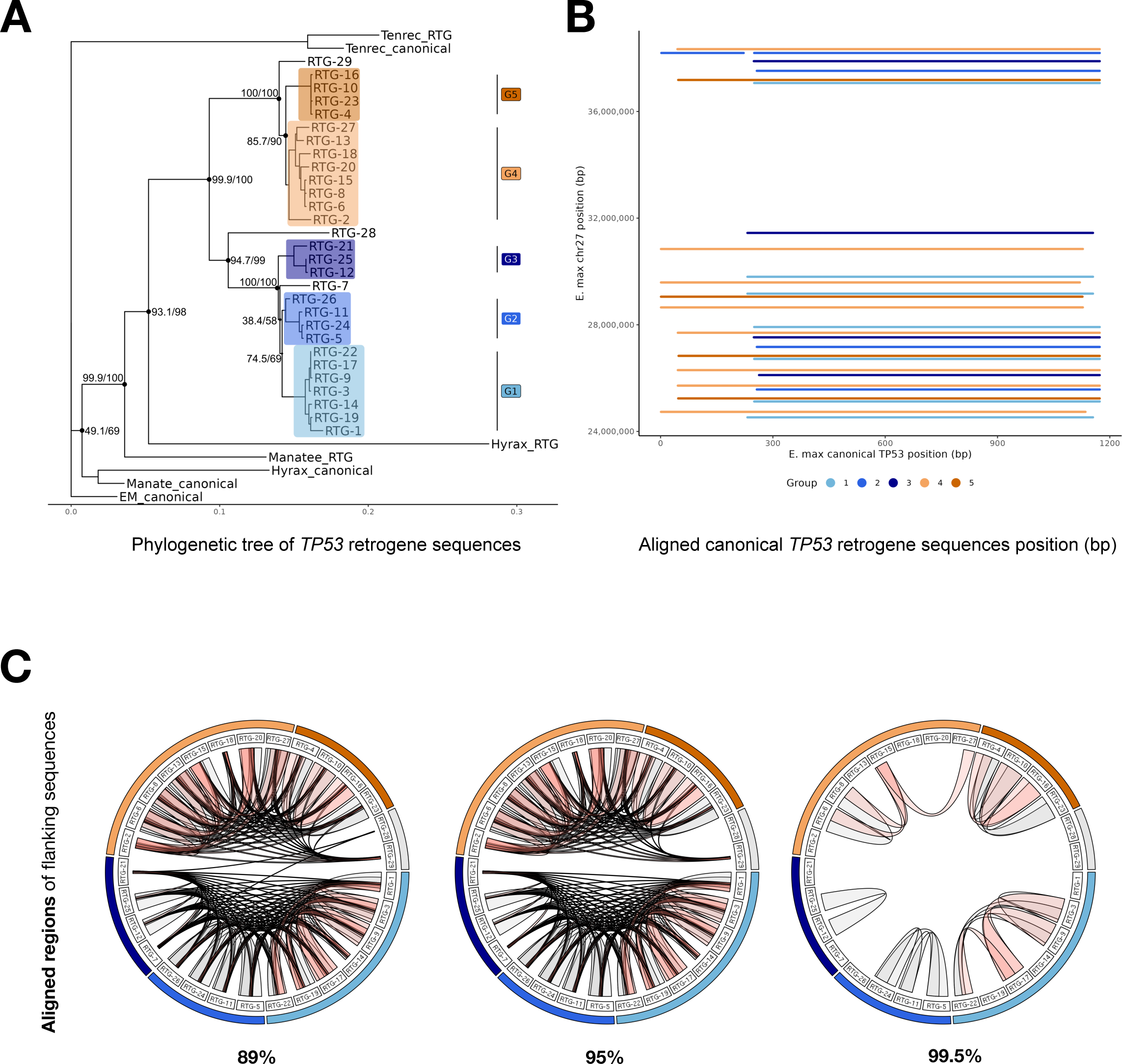
Analysis of the *TP53* RTG sequences in mEleMax1. **A:** Phylogenetic tree of the 29 mEleMax1 *TP53* RTGs. The group A and B along with their subtypes (G) are indicated in shades of blue or orange. The numbers in parentheses correspond to the SH-aLRT support (%) and ultra-fast bootstrap support (%), respectively, with higher values of both tests indicating stronger support for the branch. **B:** DotPlot and illustration of the location of p53 coding sequences of *TP53* RTG 1 to 27, identified on Chr. 27 of the Asian elephant. Blue dots represent those of group A sequences and orange dots those of group B. **C:** Circus plots illustrating the alignment among the extended regions around each *TP53* RTG (50 kb flanking sequences), filtered by similarity percentage: 89% (i), 95% (ii) and 99.5% (iii).

### Analysis of genetic elements and coding sequences

The alignments of each region flanking the *p53* RTGs (**Supplementary Table 5**) revealed sequences that share similarity, named *p53* proximal regions “PPR”. Gene and repeated sequences within each PPR were identified with BLAST v. 2.4.0 [21] and RepeatMasker v.4.0.9 [26] with default parameters **(Supplementary Table 6)**. To perform a detailed analysis of the coding sequences of the Asian Elephant *TP53* copies, we determined the putative p53 proteins, their length, and the conservation of each BOX-I domain, transactivation domains (TAD1, TAD2) and DNA Binding Domain DBD **(Figure 3B,C)**. The 29 *TP53* RTGs were grouped in three categories depending on their potential protein-coding lengths and multiple alignment of representative sequences with the canonical elephant *TP53* and with the extensively annotated human p53 (BAC16799.1) was performed **(Figure 3B,C)**. To identify genetic elements located near *TP53* RTGs, we downloaded the Repeat Masker track from the UCSC Genome Browser (//genome.uc-sc.edu) for the Asian elephant, manatee, tenrec and hyrax. Only repetitive elements within +/- 50 kb of each RTG were retained for representation. The illustration was prepared using the package Ggplot in R. MotifFinder (https://bioweb.pasteur.fr/docs/modules/seqan-library/1.3.1/CLASS_Motif_Finder.html) and BLAST v. 2.4.0 [21] were used as additional tools to validate the genetic elements and open reading frames (ORFs) in the *TP53* RTG flanking regions (integrated in **Supplementary Table 7**).

### Synteny analysis

Genes flanking the Asian elephant *TP53* RTG cluster on chr. 27 were identified from the UCSC Genome Browser (//genome.ucsc.edu). These genes were blasted to the GRCh38 human genome and the corresponding region was detected on Chr. 3. In addition, five additional genome sequences were used to identify the syntenic relationships with the analyzed *E. maximus* genome (GCF_024166365.1): the previously used *L. africana* African elephant genome (loxArf3, GCA_000001905.1); another *E. maximus* genome (GCA_033060105.1) [27]; and the following phylogenetically close species: *Procavia capensis* (Hyrax, GCA_004026925.3)*; Trichechus manatus latirostris* (Manatee, GCA_030013775.1); and *Echinops telfairi* (Tenrec, GCA_000313985.2). We compared in a pairwise manner, the chromosome-grade assembly of the Asian elephant (EleMax, GCA_024166365.1) **(Supplementary Table 1)**. The Nummer software [28] was used to detect pairwise syntenic regions between the Asian elephant and all other genomes, and alignments with more than 80% identity were selected.

## Results

### The *TP53* RTGs form two main phylogenetically distinct groups and present copy number and sequence variations in elephant species

First, to compare our results with the previous analysis in African elephant [11, 18, 20], the canonical *TP53* transcript sequence (XP_010594888.1) was aligned using BLAST against LoxAfr3 and LoxAfr4 [20]. As expected, 18 additional *TP53* RTGs hits were detected in LoxAfr3, whereas there were 18 in LoxAfr4 **(Supplementary Table 2)**. Sequences from each assembly were further aligned in pairs and the identity percentages range from 87% to 99% in LoxAfr3 and from 83% to 85% in LoxAfr4 (**Supplementary Table 3**). As already suggested [12], *TP53* RTG sequences from LoxAfr3 grouped based on their identity in two main sets, including RTGs 1-6 (Type A) and RTGs 7-18 (Type B), exhibiting >95% identity between the copies within each group **(Supplementary Table 3)**. A combined analysis of all the RTG sequences showed that LoxAfr3 Type A RTGs were highly identical (above 95%) to loxAfr4 RTGs 4,5,6 and 8, while LoxAfr3 Type B was highly identical to LoxAfr4 RTGs 7, 9-18 **(Supplementary Table 3)**. The corresponding sequences of each RTG in LoxAfr3 and LoxAfr4 were estimated based on the highest sequence identity, with values ranging from 97.5% to 100% **(Supplementary Table 3B)**. However, three LoxAfr4 RTGs (1-3) had not a clear correspondence in LoxAfr3 **(Supplementary Table 3B).** The detected variation may thus result from assembly differences generating slightly different RTG copies. As such, a one-to-one assignment of the RTGs in the two African elephant genomes could not be conclusive, in line with previous findings [11]. In addition, as these scaffold assemblies are not located at the chromosomal level, the genome organization of the RTG copies cannot be inferred.

In order to explore the retroposition and duplication events that led to the amplification of the *TP53* sequences in elephants, it is vital to determine the position of the RTGs on the genome. To that purpose, we used the Asian elephant chromosomal genome assembly (mEleMax1). By blasting the African elephant canonical *TP53* sequence against the mEleMax1 genome, 29 hits were found with 83% to 87% identity **(Supplementary Table 2C)**. The pairwise comparison of the identified RTGs between them and with those of LoxAfr3 showed that they exhibited high identity (>95%) with the two previous groups within the same species and between species, with 13 and 15 of the Asian elephant genomes belonging to Type A and Type B, respectively **(Supplementary Table 4)**. The only exception was *TP53* RTG28, which showed 84.2-89.3% identity with all other RTG copies in the two species. The phylogenetic tree of the multiple alignment of the 29 mEleMax1 *TP53* RTG CDSs using the IQ-TREE software illustrated the separation of Type A from Type B **(Figure 1A)** and also revealed several possible additional subgroups within each type (groups 1, 2 and 3 from Type A and groups 4 and 5 from Type B), which are maintained in the combined tree of all the LoxAfr3 and mEleMax1 *TP53* RTG CDSs (**Supplementary Figure 1)**. as expected by the disparity in RTG numbers, the correspondences of RTGs between species are not one-to-one and several copies from mEleMax1 exhibit similar high identity to specific RTGs from LoxAfr3 **(Supplementary Table 4)**. These results indicate that the two *TP53* RTG groups precede the divergence of African and Asian elephants, accumulating variation within each species.

We then focused on the chromosome level assembly of the Asian elephant to investigate the chromosomal positions of each RTG. *TP53* RTGs 1-27 are located in Chr. 27, while RTG 28 and 29 are at Chr 26 and 24, respectively, with RTGs 1, 2, 16-21, 28 and 29 in forward direction and 3-15 and 22-27 in reverse direction **(Supplementary Table 2C)**. Interestingly, the positioning of the *TP53* RTG types on Chr. 27 follows certain patterns, with consecutive pairs of Type A and B RTGs: e.g. RTGs 1/2, 3/4 5/6, 7/8, 9/10, 12/13, 22/23 or 26/27; and in a reversed order 16/17 and then 18/19 **(Figure 1B)**. These results suggest that the Chr. 27 *TP53* RTG cluster could have been formed by an ancestral duplication of a Type A/B pair that was followed by additional duplication events.

In order to track the duplication events, the 50-kb downstream and upstream sequences flanking each *TP53* RTG **(Supplementary Table 5)** were analysed by circus plots of increasing identity cutoff were produced to illustrate the alignments of the extended *TP53* RTG regions **(Figure 1C)**. The 99.5% identity alignments showed just the similarity between some of the copies belonging to each of the 5 subtypes observed from the tree, while the 95% identity alignments showed the division of Types A and B **(Figure 1C)**. The alignments of 89% identity illustrated some additional connections between the RTGs within the two groups **(Figure 1C)**. These results support some of the phylogenetic relationships indicated by the tree **(Figure 1B)**, such as that RTG 29 (Chr. 24) is closest to Group B. In addition, ie the sequence around RTG 28 is 89% identical to that of RTG 7 and RTG 21 **(Figure 1C)**, suggesting common ancestry with Group A. Together, these results show that the evolution of the duplicated sequences involving p53 are extended around p53 by several kb, and during their expansion, two main types and five groups were formed in three different chromosomes.

### Genetic elements in proximity to *TP53* RTGs and syntenic analysis of the duplicated regions

We next set up to further analyse the 29 extended regions that comprise the *TP53*-like sequences to investigate the genetic elements in proximity to these sequences and try to understand better the evolution of the duplications. First, we performed a detailed characterization of the repetitive element content of the 100 kb sequences flanking the 29 *TP53* RTGs using Repeat Masker, which was in line with previous studies [12] **(Supplementary Table 6)**. However, several patterns formed by combinations of different types of repetitive elements were detected upstream and downstream of each *TP53* RTG that match well the five subtypes of the phylogenetic tree **(Figure 2A)**, further confirming that these elements were duplicated along with the *TP53* RTGs. The subtypes within each main group were clearly separated by large differences, such as the downstream sequences between subtype 4 and 5, or smaller deletions and insertions of repetitive sequences, like those differentiating subtype 1 from 2 and 3, which are more similar to each other, and subtype 2 and 3 between themselves **(Figure 2A).** This confirmed that RTG 7 belongs to sub-type 3, which was not well resolved from the *TP53* RTG tree. In addition, in some RTG copies there are specific extra insertions that support their common origin, like those upstream and downstream of RTG 14 and 19 **(Figure 2A)**. Therefore, the repetitive element analysis provides additional information on the relationships between some of these RTG copies and how they were generated. For example, there are four pairs of closely-related RTGs of subtype 1 (3, 9, 17 and 22) and 5 (4, 10, 16 and 23) **(Figure 1A**; **Figure 2A)** that are always located together in the genome, suggesting that they were originated by independent duplication events. Similarly, the repetitive element pattern indicates a close relationship of RTG 29 with all the others Group B RTGs, whereas RTG 28 shows a quite different repetitive element content, but the region surrounding the *TP53* RTG is very similar to those of Group A **(Figure 2; Supplementary Table 7)**.

**Figure 2.**
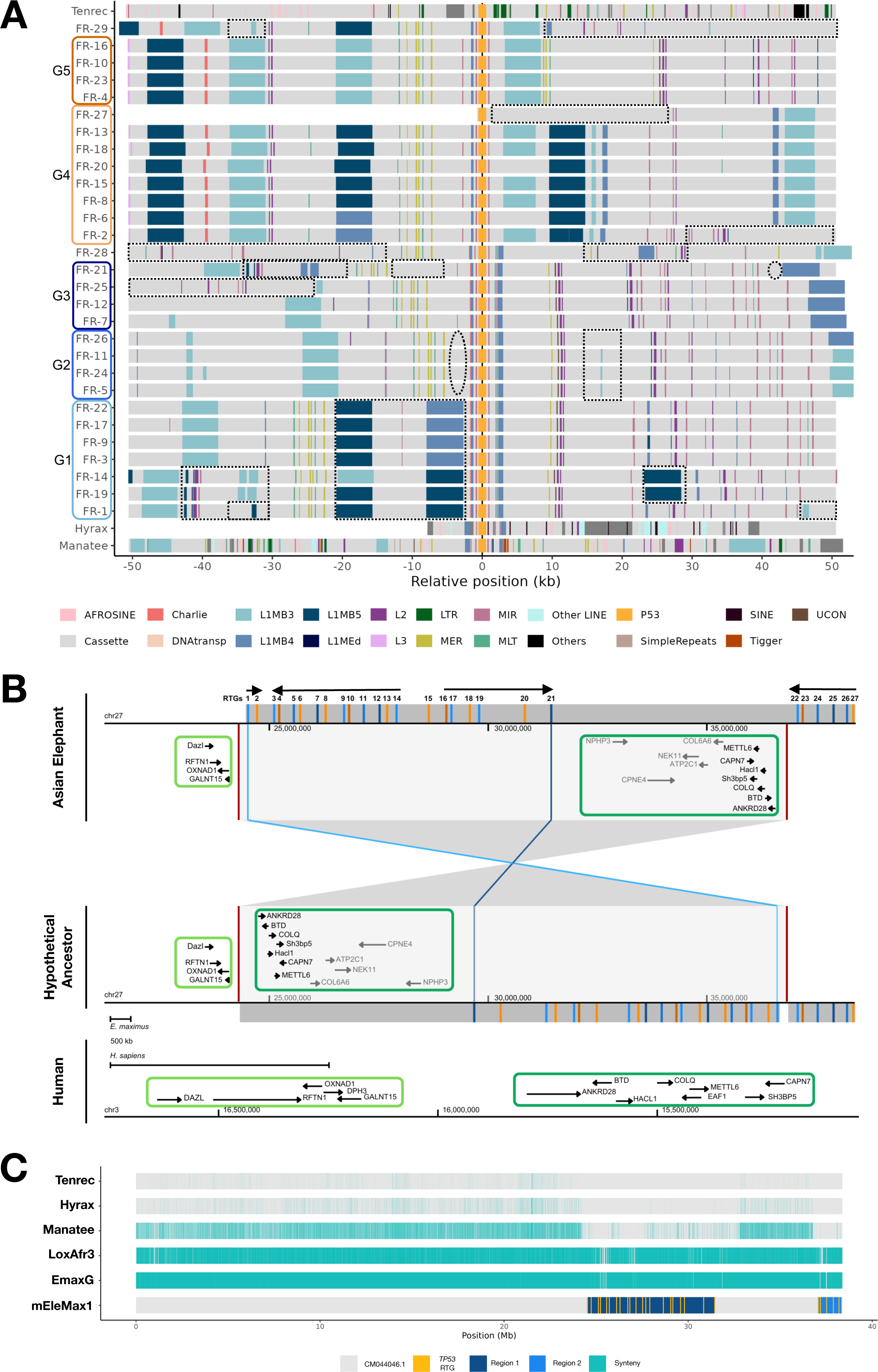
Illustration of genetic elements and repetitive elements flanking the *TP53* RTGs in the Asian elephant and in related species. **A:** Plot showing the flanking region (FR) including the repetitive elements in proximity (+/- 50 kb) to each *TP53* RTGs (coloured orange). Five types of repetitive element patterns are observed in the elephant and three diverse patterns in related species, in line with the phylogenetic analysis of the *TP53* RTG sequences. Sequences coloured in white are not available in the respective assemblies. Possible insertions are indicated with dotted rectangles and putative deletions are indicated by dotted circles. Collectively, the following repetitive elements were detected **(Supplementary Table 6)**: MIRb, MIRc, MER20, MIR, MIR3, MLT1A, MLT1B, MER5A, MER5B, MER3, MER20B, L1MEd, L1MB4, L1MB5, GA-rich, (A)n, (T)n, L1MB4, UCON2, L2a, Charlie4a, Charlie4. Tigger20a elements were also identified (not shown for simplicity). **B:** Comparative analysis of the region on Chr 27 of the Asian elephant including 27 *TP53* RTGs, with the corresponding region on human Chr3 in human. The orientation of the copies is indicated in black arrows. The correspondence is based on the conservation of two groups of protein-coding genes in proximity to the amplified *TP53* RTGs in the elephant. The hypothetical inversion of RTGs 1 and 22 in the elephant is illustrated. **C:** Illustration of syntenic analysis of the *TP53* RTG cluster found on Chr. 27 of the Asian elephant. Green-shaded areas denote the syntenic regions between the Asian elephant and the rest of the genomes. Regions in blue, that contain the *TP53* RTGs (in yellow), are exclusively found in elephant genomes. The chromosome-grade assembly of the Asian elephant (mEleMax1, GCA_024166365.1) was compared against the following genomes: the EmaxG (GCA_033060105.1) and the scaffold-grade assemblies from the African elephant (LoxAfr3, GCA_000001905.1), the Hyrax (GCA_004026925.3), the Manatee (GCA_030013775.1) and the Tenrec (GCA_000313985.2).

To identify additional genetic elements further analyse the extended regions in proximity to each *p53* RTG, we then performed BLAST analysis using each extended region of 50kb flanks as query. In particular, consistent with previous studies [12], a repeated region, here annotated as p53-proximal-repeat (PPR), was detected in proximity to all *TP53* RTGs. The length of the aligned PPR sequences varies from 4 kb (e.g. in RTG 12) to approximately 7 kb long (e.g. in RTG 14) and contain ORFs that show high similarity with parts of LINE1 and RT_nLTR elements (**Supplementary Table 6)**. Blast analysis of the PPR region with available protein sequences showed no homology with other coding genes, except some putative proteins from Asian elephant, such as receptor tyrosine-protein kinase erbB-4 isoform X3 (XP_049744894.1), which seems to be just mistake in the genome automatic annotation **(Supplementary Table 7)**. Therefore, no additional gene candidates that might be involved in the maintenance of these duplicated sequences were found.

### Syntenic analysis of the *TP53* RTG regions

To investigate in more detail the chromosomal positions of the *TP53* sequences evolved in the elephant lineage, we performed a synteny analysis at different evolutionary distances. First, we compared in a pairwise manner, the chromosome assembly of the Asian elephant with the scaffold assemblies from the African elephant and three phylogenetically related species: hyrax, manatee and tenrec. We also tested the synteny with another Asian elephant genome that was released at the time of the analysis (EmaxG) [27], which showed a very similar organization **(Figure 2C).** Results focusing on the Asian elephant chromosome 27, showed that the whole region comprising the 27 duplicated *TP53* RTG copies are elephant-specific and absent in other phylogenetically close species, in line with previous findings employing other genome sequences [12, 17] **(Figure 2C)**. In addition, the absence in this region of the possible phylogenetically related *TP53* RTGs in hyrax and manatee **(Figure 1A)**. suggests that it is not the location of the original insertion event **(Figure 2C)**.

Next, we identified sets of protein-coding genes that are located upstream and down-stream of the *TP53* RTG Chr. 27 clusters in mEleMax1 and compared them with those in humans. We focused on two distinct regions, depending on the proximity of the RTG copies, including 1 to 21 and 22 to 27, which are relatively close to Chr. 27 telomere **(Figure 2B)**. Thereof, the distal genes to the first region include *SATB1, TBC1D5, PLCL2, DazI, RFTN1, OXNAD1,* and *GALNT15*; while the genes between both regions are *Olfr917, Gm10310, NPHP3, ACAD11, ACKR4, DNA-JC14, ACPP, CPNE4, MRPL3, NUDT16, NEK11, ATP2C1, COL6A6, PIK3R4, COL6A5, CAPN7, SH3bp5, METTL6, EAF1, COLQ, Hacl1, BTD* and *ANKRD28*. Interestingly, in the human genome the first and second block of genes are close together, with the latter being in reverse order and orientation compared to the Asian elephant genome **(Figure 2B)**. In particular, the distance between the *ANKRD28* and *GALNT15* genes, which mark the limits of the two blocks, is 12.5 Mb in the Asian elephant and 377 kb in humans **(Figure 2B)**. Thus, this supports the insertion and duplication of the *TP53* RTGs between *ANKRD28* and *GALNT15* to create the elephant Chr. 27 cluster, in line with the the synteny analysis in closer species, which then got separated as a result of an inversion.

### The duplicated copies of truncated p53 sequences partially retain putative functional domains

Finally, to gain additional insights on the potential activity of the Asian elephant *TP53* RTGs, we analysed the conservation of the p53 coding sequences. The length of the putative encoded proteins varies from 26 aa to 212 aa and can be grouped in three main classes: 10 RTGs encode truncated p53 sequences of about 200 aa; 12 RTGs of about 150 aa; and 7 RTGs of up to 79 aa **(Figure 3B; Supplementary Table 8)**. All sequences contain the BOX-I motif of the transactivation domains (TAD-I and TAD-II), which was assigned based on its identity with the corresponding BOX-I type motifs in the African elephant [14, 15]. Moreover, they include partial putative DNA binding domains (DBD) sequences, i.e. 16% in the RTGs of about 150 aa and 60% in the RTGs of about 200 aa, with several differences compared to the canonical p53. Analysis of the putative p53-coding sequences also showed that 23 of the *TP53* RTGs contain the start codon ATG, 2 contain CTG and 2 do not include a start codon **(Supplementary Table 8)**. A methionine coding ATG codon is present in RTGs 3, 7, 9, 17, 21, 22, and 26 encoding for 200 aa long putative p53 sequences, which also include a variety of BOX-I motifs and maintain 60% of the corresponding DBD length. In addition, all RTGs of about 150 aa, maintaining 16% of the DBD length, have start codons, with the exception of RTG 12 and 25. Conversely, RTGs 4, 10, 16, and 23 have start codons and contain the TADI and TADII, but lack the DBD **(Figure 3B,C).** Therefore, there are two main pools of putative p53 truncated proteins, one of less than 150 aa expressing the Type B BOX-I (QETFSYLGKLLPEKLV) and either none or only 16% in length, of the DBD length, and a second one of >200 aa that expresses 60% of the DBD and either the Type E (SQETFSYLGKLLSEKLV) or Type F (SQGTFSYLGKLLPEKLV) BOX-I motif. An example of the alignment of one RTG of each set is shown in **Figure 3C**. Actually, this classification fits closely with that of RTG types based on the tree and repetitive elements, with Type 1 encoding longer proteins with L120, Type 2 encoding shorter proteins, except RTG 26 (longer protein with L120), Type 3 encoding longer (RTG 7 and 21) or intermediate (RTG 12 and 25) proteins with L120, Type 4 encoding an intermediate protein with the original K120, and Type 5 encoding just shorter proteins **(Figure 3B)**. Interestingly, the shorter DBD (e.g. RTG 27) are well conserved compared to the canonical elephant p53, while the longer DBDs (e.g. RTG 17) are poorly conserved. The RTG 21 that retains 60% of the DBD, corresponds to *p53* RTG9 of the African elephant, which was shown to induce apoptosis [13]. Our analysis suggests a variable conservation for the duplicated *TP53* RTGs, notably of the DBDs, pointing towards diverse functions. In line with previous data on the MDM2/p53^BOXI^ interactions [14, 15], this variation indicates variable potential to trans-activate downstream targets, forming functional pools that cooperatively participate in DNA binding activities [29].

**Figure 3.**
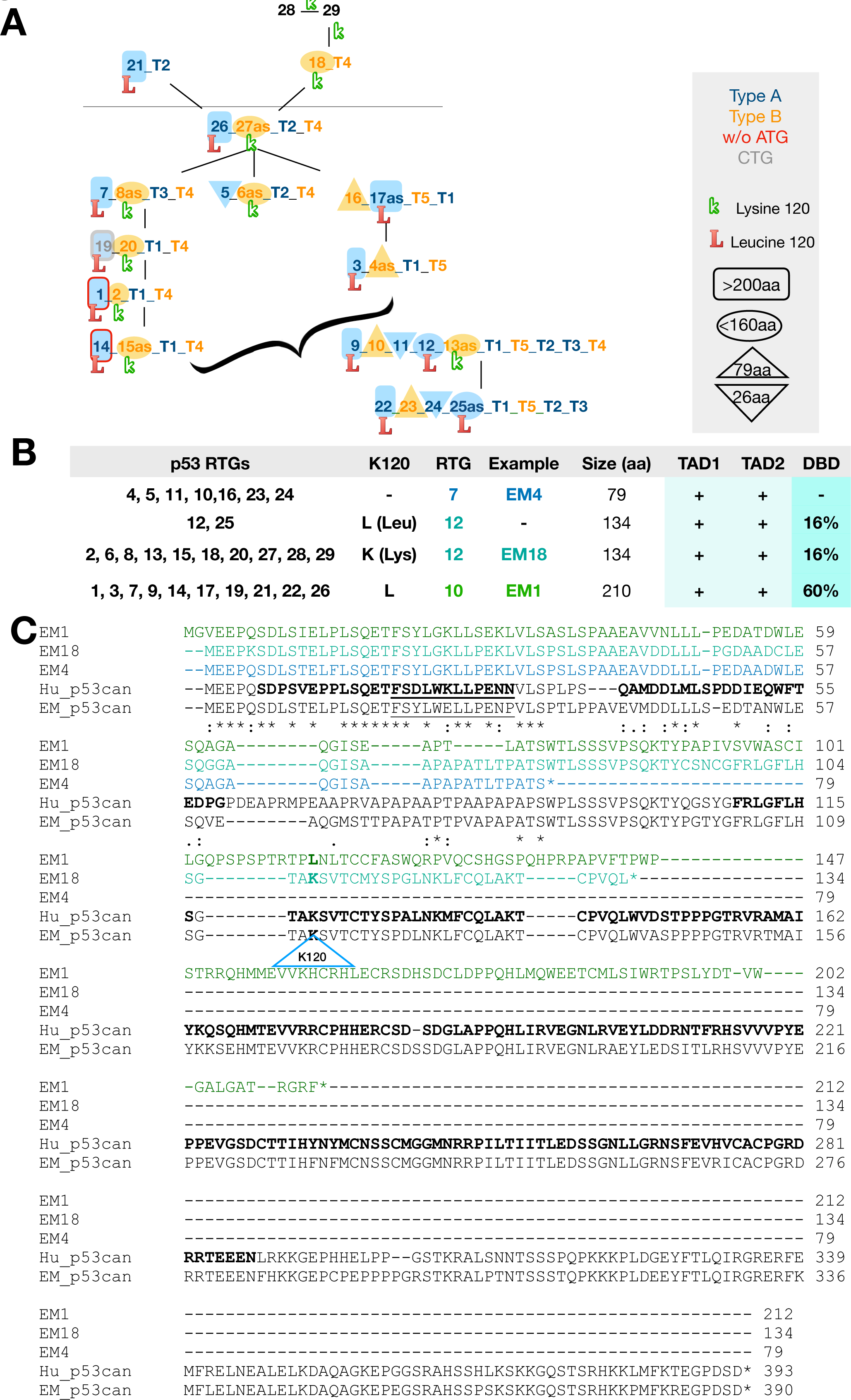
Characterisation and evolution of *TP53* RTGs. **A:** Model proposing the series of duplications leading to the amplification of *TP53* RTGs, their types and protein sizes in the Asian elephant. **B:** Summary and analysis of the p53 sequences encoded by *TP53* RTGs. The transactivation domains TAD and the partial DBD encoded by *TP53* RTGs are aligned with the annotated human p53 sequence. Based on the length of the putative CDS of each *TP53* RTG and the retaining of the TAD and DBD domains, three groups are formed (∼79 aa, marked in blue; ∼134 aa, marked in light blue, and ∼210 aa, marked in green). **C:** Sequence alignment of different putative *p53* proteinsRTG encoded sequences represent the three main types of different lengths in Asian elephant (RTG 4: 79 aa; RTG 18: 134 aa; RTG 1: 210 aa). The K120L substitution in RTG 1, that is crucial for the acetylation of p53 in human, is indicated. The sequences encoding for the TADI and TADII and for the corresponding putative DBDs are highlighted in bold font.

## Discussion

### The investigation of the unique elephant multi-*TP53* system necessitates accurate genomic assemblies across elephant species

Throughout evolution, the ancestral *TP53/63/73* gene was involved in development and gave rise to three distinct genes in vertebrates and mammals with roles in DNA repair and numerous other functions (e.g. in humans, p53 has roles in cancer and DNA damage-induced cellular apoptosis, while p63 and p73 have roles in development). Human p53 incorporates its multiple functions into a single sequence that is tightly-regulated via post translational modifications (PTMs) [3], but several truncated p53 spliced isoforms have also been described [30–32]. Elephants, on the other hand, maintain multiple truncated *p53* RTGs [12, 33, 34]. Here, we extend this knowledge by mapping them on a chromosome-level genome assembly of the Asian elephant, a phylogenetic classification of both the *p53* sequences and the flanking REs, and a detailed evolutionary analysis of the evolution of these copies and how they originated. Even though these copies are partial and originate from retrotransposition that normally results in non-functional pseudogenes, the variety of p53-like proteins helps in understanding evolutionary mechanistic aspects of the regulation of p53, both at the structural and the signaling levels. Evidence already suggest that these copies induce apoptosis via the p53 signalling pathway resulting in an enhanced sensitivity of elephant cells to DNA damage (compared to the closest living relatives) [35].

However, due to limited coverage or read lengths, the available assemblies may be suboptimal, or the detected copies may also be polymorphic in the population, which precludes having a good idea of this gene system. Regarding the Asian and African elephant, in three earlier studies [11, 12, 17], *TP53* RTGs were examined using scaffold-level genomes (LoxAfr3, LoxAfr4, or draft Asian assemblies), supplemented with capillary re-sequencing and whole-genome Illumina data [11] or Illumina short-read data from Asian elephants [12]. Here, we instead use the newer chromosome-level mEleMax1 assembly for the Asian elephant, alongside with comparisons to LoxAfr3 and LoxAfr4, providing a clearer view of location of *p53* RTGs on chromosomes and the variation among them. In line with this, based on read depth, African elephants were estimated to have ∼18–23 *TP53* RTG copies and Asian elephants 10 to 37 copies [17]. In fact, capillary sequencing on single elephant individuals resulted in clustered but not identical *TP53* RTG sequences compared to data available in previous annotations, showing that there exists considerable sequence variation [11]. Consistent with previous results, we identified 18-19 copies in the African elephant and 29 copies in the Asian elephant. Moreover, based on the phylogenetic tree of the CDS and the flanking repetitive elements, we showed that, besides the two previously described main groups, the *TP53* RTGs can be further categorised in three subtypes for group A (1, 2 and 3) and two for group B (4 and 5). Moreover, building on Sulak et al. (2016), who reported that *TP53* RTGs expanded via a single retrotransposition event followed by segmental duplications [12], we demonstrate here that each *TP53* RTG is flanked by subtype-specific TE combinations (**Figure 2A**). The identified repetitive elements are in line with previous studies on the African elephant. MER5A/MIRc, RTE1/MLT1J1, RTE1_LA, Charlie4, L1M-B4, MIR3 were previously identified upstream of *TP53* RTG; and MIR, MLT1X_LA, L1MB4/ MIR3 downstream of *TP53* RTG [12]. This supports a stepwise duplication model in which the repetitive elements were duplicated along with the *TP53* RTG copies, and provides finer resolution of elephant-specific genomic architecture **(Figure 2B**, **Figure 3A)**. As already documented [12], comparative analysis with phylogenetically related species indicates that elephant-specific RTG expansion is absent in Manatee, Hyrax, and Tenrec, supporting the hypothesis that the observed duplication events occurred after the elephant lineage diverged. However, by focusing on both the *TP53* sequence and the flanking repetitive elements, we found that heir patterns in Manatee and Hyrax are more similar to elephant Group A than Group B RTGs **(Figure 2A).** Tenrec on the other hand shows a distinct pattern and identity of repetitive elements flanking the *TP53* RTG and it is missing L1MB3/4/5 elements that are abundant in the elephant and present in Hyrax and Manatee **(Figure 2A)**, which confirms previous observations indicating that Tenrec *TP53* RTG results form an independent retrotransposition event [12].

### A step-wise amplification of *TP53* copies

By integrating (i) the phylogeny of *TP53* RTGs, (ii) the alignment of their flanking sequences and the repetitive element patterns, and (iii) the synteny analyses in closely related species and humans, we propose a possible stepwise model of *TP53* RTG amplification in the Asian elephant **(Figure 3A)**. As already mentioned [12], first, a single ancestral retrotransposition gave rise to the original *TP53* RTG insertion, which likely resembled more those of Group A, and included the common repetitive elements that form the PPR of all RTGs. The lack of synteny of the Chr. 27 RTG cluster with the single copies in hyrax and manatee and the similarity of the closest repetitive elements suggests that the original insertion probably corresponds to RTG 28 in Chr. 26. Next, this ancestral PPR region was probably duplicated at the end of Chr. 27 and the two *TP53* RTG copies diverged for some time, giving rise to the main RTG groups A and B. At some point, RTG 28 was likely copied also into the end of Chr. 27, during which the canonical Lys in position 120 was substituted by a Leu, and the merged pair of Group A and B RTGs were duplicated again, generating the four ancestral copies that resulted in the different *TP53* subtypes: T1, T2/3, T4 and T5. Finally, additional sequential duplications of one or several of these genes together, followed by other rearrangements of the region favored by the sequence similarity, generated the current distribution in the available Asian elephant genome, with mainly alternating Group A and B RTGs **(Figure 2A)**. For example, there are four representatives of a possible cluster formed by subtypes T1, T5 and T2, most of which show high sequence similarity, which in some cases occur together with a subtype T4 RTG (3-4-5-6 and 17-16-26-27, although the two parts of this cluster are separated in two regions of Chr. 27) or a subtype T3 RTG (9-10-11-12 and 22-23-24-25. Also, other pairs of subtype T3 and T4 RTGs that could have been generated by duplication are 7-8 and 12-13. As mentioned in the results, synteny with humans suggests that a large inversion in the elephant lineage moved a large part of the *TP53* RTG cluster to a more proximal position. In addition, assuming that all the copies had originally the same orientation, another inversion could have inverted the block from RTG 3 to 15 with respect to the rest of the copies.). According to this model, RTG 29 probably resulted from an independent duplication of a subtype 4 copy, including most of the upstream and a small part of the downstream flanking region.

This model explains the observed phylogenetic structure of the *TP53* RTG copies and provides a framework for future evolutionary and functional studies. Nevertheless, the complexity of the region emphasizes the need for additional sequencing data and chromosomal-level assemblies, that can be obtained with kb-long reads targeting the consecutive elephant *TP53* copies on Chr. 27. Such data could be enriched by adaptive sampling for long reads from Oxford Nanopore Technologies (ONT), providing information on the potential polymorphic nature of RTG copy numbers within and between elephant species and some crucial intermediate steps in the generation of the observed diversity of this gene family. Similarly, tracing the evolution of the *TP53* RTG copies in extinct species of the elephant lineage, such as mammoths [36, 37] or dwarf elephants [38, 39] could help validate the proposed model on the *TP53* RTG duplications and their functional implications [40, 41].

### The duplicated p53 copies retain putative TAD and DBD functions

While this study provides a detailed map of *TP53* RTGs in elephants, their expansion should not be interpreted solely as an adaptation for cancer suppression. As recently argued [42], many of the mechanisms attributed to cancer resistance are more parsimoniously explained as part of broader somatic maintenance programs, with cancer suppression emerging as a secondary benefit. In line with this, it has been hypothesised that *TP53* RTGs may uptake roles in the preservation of the germline. Because elephant testicles remain undescended, sperm production occurs at elevated body temperatures, which may place additional stress on DNA replication and repair in the germline [19]. The current detailed analysis therefore provides a foundation for functional studies that can disentangle the roles of elephant *TP53* RTGs in general stress response and tissue maintenance, as well as their possible contribution to reduced cancer prevalence.

At the molecular level, the elephant *TP53* RTGs retain modified sequences of the entire TAD and partial ones for the DBD **(Figure 3B)**, suggesting potential functional diversity. While detailed functional validation is required, this structural variation likely influences p53 signaling and evolutionary adaptation.

**(i)** The BOX-I motif at the p53 TAD (N’ terminal) is key for the interaction and regulation by MDM2 [3, 4, 6]. Even though it is well-conserved throughout evolution, variations were shown to modify its structure and interaction interface with MDM2 [7, 8, 14]. It is worth mentioning that the N’ termini (BOX-I motifs) of all *TP53* RTGs encoded putative proteins were shown to adopt more stable conformations compared to human p53 [15]. However, all elephant *TP53* RTGs encode a W to G substitution of the FxxxWxxL motif [12], which was previously shown to contribute to a decreased binding capacity of p53 with MDM2 [14] that is dynamically affected by temperature [15]. The double mutant p53^25,26^ (L22Q, W23S) in mice is severely compromised for the capacity to activate most p53 targets, but it is fully active on a subset of them (e.g. *BAX*). Displaying a selective transactivation function, p53^25,26^ induces mid-gestational embryonic lethality [43]. Interestingly, while p53 TAD is required to suppress expression of SINEs and LINE-1 elements [44] the p53^25,26^ mutant phenocopies the ability of wild type p53 to maintain suppression of B1 or B2 SINEs expression, when treated with the DNA demethylating agent 5-Azacytidine (5-Aza). As 5-Aza treatment induces expression of repetitive elements in p53-deficient cells, p53 is proposed to be involved in the DNA methylation machinery to silence repetitive elements [45]. In line with previous studies showing that p53 mitigates oncogenic disease in part by restricting transposon mobility through its involvement in the DNA methylation machinery [45, 46], experiments testing the capacity of the truncated elephant *TP53* RTG encoded proteins to control the methylation patterns are anticipated to reveal potential roles of these sequences in development or cancer.
**(ii)** The DBD, folded by the C’ terminal of p53, interacts with target DNA sequences, (p53REs, following the 5’ RRRCWWGYYY ‘3 sequence), enabling stress-dependent transactivation of downstream genes, such as the p21encoding gene *CDKN1A* or apoptotic targets like *BAX* [47]. In elephants, several *TP53* RTGs retain only partial DBDs, indicating that these truncated copies most likely do not retain the structural features of canonical p53 **(Figure 3B,C)**. However, 10 *TP53* RTGs retain 60% of the DBD. Notably, none of the elephant truncated p53 putative proteins retains the C’ terminal region of the canonical protein, which in humans is known to regulate pro-apoptotic structure-selective DNA binding. In fact, mutations on the C’ terminus were previously shown to lead to altered specificity to local DNA structures, playing critical roles in human tumor progression. The apparent lack of the p53 C’ terminus in the elephant’s *TP53* RTG sequences documented here may thus lead to increased intrinsic disorder in the DBDs with functional consequences.

It is worth mentioning that while p53 is known to regulate gene expression as a tetramer, monomeric p53 proteins have also been shown to regulate cell fate by binding to the p53RE DNA sequences. In particular, p53 monomers were suggested to overcome the dominant negative effect of mutant forms of p53, which disrupt the functional p53 tetramers, leading to cancer [48]. In line with this, several studies point towards alternative functions of the truncated *TP53* RTGs that do not involve transactivation of p53 targets. Indeed, the African elephant TP53 RTG9, which encodes a truncated protein that only retains half of the DBD and lacks the C’ terminal oligomerization domain that is required for the tetramerization of p53, recently was shown to bind to the Tid1 mitochondrial chaperone in response to cellular stress and induce apoptosis [13]. This seems to correspond to RTG 26 in the Asian elephant and has high similarity with other Group A *TP53* RTGs. The structure of these elephant *TP53* RTGs resemble that of the human p53Ψ, a transcriptionally inactive p53 isoform alternatively 3′ spliced in intron 6 that is found in cancers. p53Ψ also has an incomplete DBD and misses the entire C’ terminal oligomerization domain. However, it uptakes alternative functions reprogramming cells towards a metastatic-like state by modulating cell survival, epithelial-to-mesenchymal transition, invasion and metastasis [49, 50]. The structural and functional diversity of the elephant *TP53* RTGs could therefore serve as natural models for functional diversification of p53, providing insights into selective p53 activation (eg preventing MDM2-mediated inhibition of the canonical p53 [14]) and offering potential strategies to stabilize or reactivate p53 in human cancer therapies. The design of modified DBDs (e.g, extended N’ or C’ termini) is proposed for translational applications promoting a selective trans-activation of downstream genes [51]. For example, designed chimeric transcription factors composed of zinc finger peptides (Zip) and p53 DBD, containing N’ and C’ terminal extensions of the DBD, increased selectively the transcription of *CDKN1A* by up to 800-fold [51]. As such, by modifying the DBD of p53, a selective activation of a set of downstream targets and the corresponding signalling pathways is achieved. Thus, in the elephant p53 model, the presented data on the role of the *TP53* RTGs expressing partial DBDs advance previous findings exploring structural implications of the p53 putative protein pools [14].[52–54]. In line with the structural evidence, stress-dependent employment of functional *p53* isoforms in human, deriving from alternative splicing or copy number variations is well documented for the *p53* gene [32, 62, 63].

Here we sought to determine and characterise the *p53* RTG copies on the genome of the Asian elephant. Taking the characterisation one step further we annotated and grouped the lengths of each DBDs. and run alpha fold models simulating their 3D structures based on a model of human p53 DBD bound on the response element of p21 (PDB 3TS8). These models reveal the topological alignment of the truncated p53 RTGs with the canonical and in relation with the bound response element. Even though it is unlikely that *TP53* RTG encoded proteins lacking the oligomerization domain can transactivate p53REs, partial DBDs may retain functions via binding other proteins, such as chaperons at the mitochondria [13], while the BOX-I motifs at the N’ terminals can contribute to the activation of the canonical p53 by competing with its binding to MDM2 (p53’s main negative regulator) [14, 15]. However, to accurately determine those structures and validate their functions, further *in silico* and *in vitro* investigations are needed. As discussed, this can be true with at least two aspects of the truncated *p53* RTGs: (i) modified BOX-I motif-carrying *p53* RTGs can target the MDM2/p53 interaction to activate the canonical p53; and (ii) truncated *p53* RTGs with partial DBD, which retain the capacity to bind other proteins (eg chaperons at the mitochondria), induce transcription-independent apoptosis at the mitochondria [13]; resembling to human P53Ψ isoform [49]. To that respect, an experimental investigation addressing whether and which of these truncated *TP53* RTGs have retained or acquired functional activities, is highly anticipated and relevant to biomimetic translational applications.

### Conclusions

In conclusion, we mapped 29 *TP53* RTGs on the Asian elephant genome, classifying them into two main groups and five subtypes with specific flanking TEs. While *TP53* RTGs in Asian elephants were previously reported [11, 12, 17], this study provides the first precise mapping using a chromosome-level assembly and how they have been originated. Compared with the 18-19 copies in African elephants and 10–37 RTGs in other Asian elephant genomes, these results indicate variable copy numbers and paralogue diversity. Synteny analysis with the human genome shows that RTGs 1–27 are organized in two separated gene clusters on chromosome 27, with an inversion between RTG1–21 and RTG22–27 **(Figure 2B)**. We also present a model indicating that the initial retrotransposed p53 copy gave rise to early RTGs on chr27 (RTG21, RTG18) and later additional rounds of duplication forming paired and extended regions (e.g., RTG1–2, 9–13, 22–25) on chr27, while additional copies emerged on chr26 and chr24 (RTG28, RTG29) **(Figure 3A)**. By describing in detail the genomic architecture and the sequence diversity, which could be linked to protein structural diversity, as shown previously [14], this work provides a framework for perspective functional studies *in vitro* and in cells.

## Supplementary Tables

**Supplementary Table 1.** Table summarising the genomes used in this study.

**Supplementary Table 2.** Blast alignment of the canonical *TP53* transcript sequence (XP_010594888.1) with LoxAfr3 (**A**) and LoxAfr4 (**B**) or mEleMax1 (**C**) genomes. List of 18-19 hits of high identity (>80%) are detected in the African elephant genomes and 29 in the Asian elephant genome. **D:** Table summarising the putative protein sequences of each *TP53* RTG identified in the Asian elephant.

**Supplementary Table 3.** Comparison of putative p53 coding sequences in *TP53* RTGs of LoxAfr3 and LoxAfr4 genome assemblies, performed with DotPlot. **A:** Table summarising the pair-wise alignments of the *TP53* blast hits from LoxAfr3 and LoxAfr4 genome assemblies. Colouring illustrates the identity (teal, 90%-95%; blue, 95%-98%; green, above 98%). Two main grops are formed based on the identity %. **B:** Table summarising the corresponding (most identical) *TP53* RTG sequence hits in either LoxAfr3 and LoxAfr4.

**Supplementary Table 4.** Comparison of putative p53 coding sequences in *TP53* RTGs of LoxAfr3 and mEleMax1 genome assemblies by pairwise alignments of the *TP53* blast hits, performed with DotPlot.. Colouring illustrates the identity (teal, 90%-95%; blue, 95%-98%; green, above 98%). Two main groups are formed based on the identity.

**Supplementary Table 5.** Table summarising the coordinates of the extended regions (extended by 50 kb), flanking each *TP53* RTGs in the Asian elephant genome.

**Supplementary Table 6.** Table summarising the repeat masker results and the coordinates of the repetitive elements flanking (100 kb) the *TP53* RTGs of the Asian elephant genome.

**Supplementary Table 7.** Table summarising the coordinates of the annotated genes (*TP53, erb-B4, GNB1L*) and repetitive elements (*RT_nLTR*) in in the 50 kb flanking regions of each *p53* RTG (PPRs).

**Supplementary Table 8.** Table summarising the amino-acidic length, the BOX-I type and the translation capacity of the p53 putative proteins in EleMax.

## Acknowledgments and funding information

This work was funded by by research grant PID2022-137615OB-I00 to MC and PID2023-146193OB-I00 to MAS funded by the Agencia Estatal de Investigación of the Ministerio de Ciencia, Innovación y Universidades (MICIU/AEI/10.13039/501100011033, Spain) and the European Regional Development Fund (ERDF, EU) and the Save The Elephants association (STE), a UK Charity (registration 118804). KK was supported by a Maria Zambrano Grant from the MICIU (Spain) funded by the European Union-NextGenerationEU to KK.

## Author contributions

KK conceptualised the study, performed analyses and wrote the paper, EC performed technical analyses and prepared figures, MP contributed to technical analysis, RF and FV conceptualised the study, MC conceptualised the study, contributed to analysis and wrote the paper. All authors reviewed the paper.

## Conflict of interest

The authors declare no conflict of interest.

## Data Availability Statement

The data that support the findings of this study are available in NCBI at https://www.ncbi.nlm.nih.gov. These data were derived from the following resources available in the public domain: - mElemax1, https://www.ncbi.nlm.nih.gov/datasets/genome/GCF_024166365.1-loxAfr3.0, https://www.ncbi.nlm.nih.gov/datasets/genome/GCF_000001905.1/-EmaxG-LGv1.0, https://www.ncbi.nlm.nih.gov/datasets/genome/GCA_033060105.1/-ProCapCap_v3_BIUU_UCD, https://www.ncbi.nlm.nih.gov/datasets/genome/GCA_004026925.3/-Trichechusmanatuslatirostris, https://www.ncbi.nlm.nih.gov/datasets/genome/GCA_030013775.1/-Echinopstelfairi, https://www.ncbi.nlm.nih.gov/datasets/genome/GCA_000313985.2/

## Supplementary Figures

**Supplementary Figure 1.**
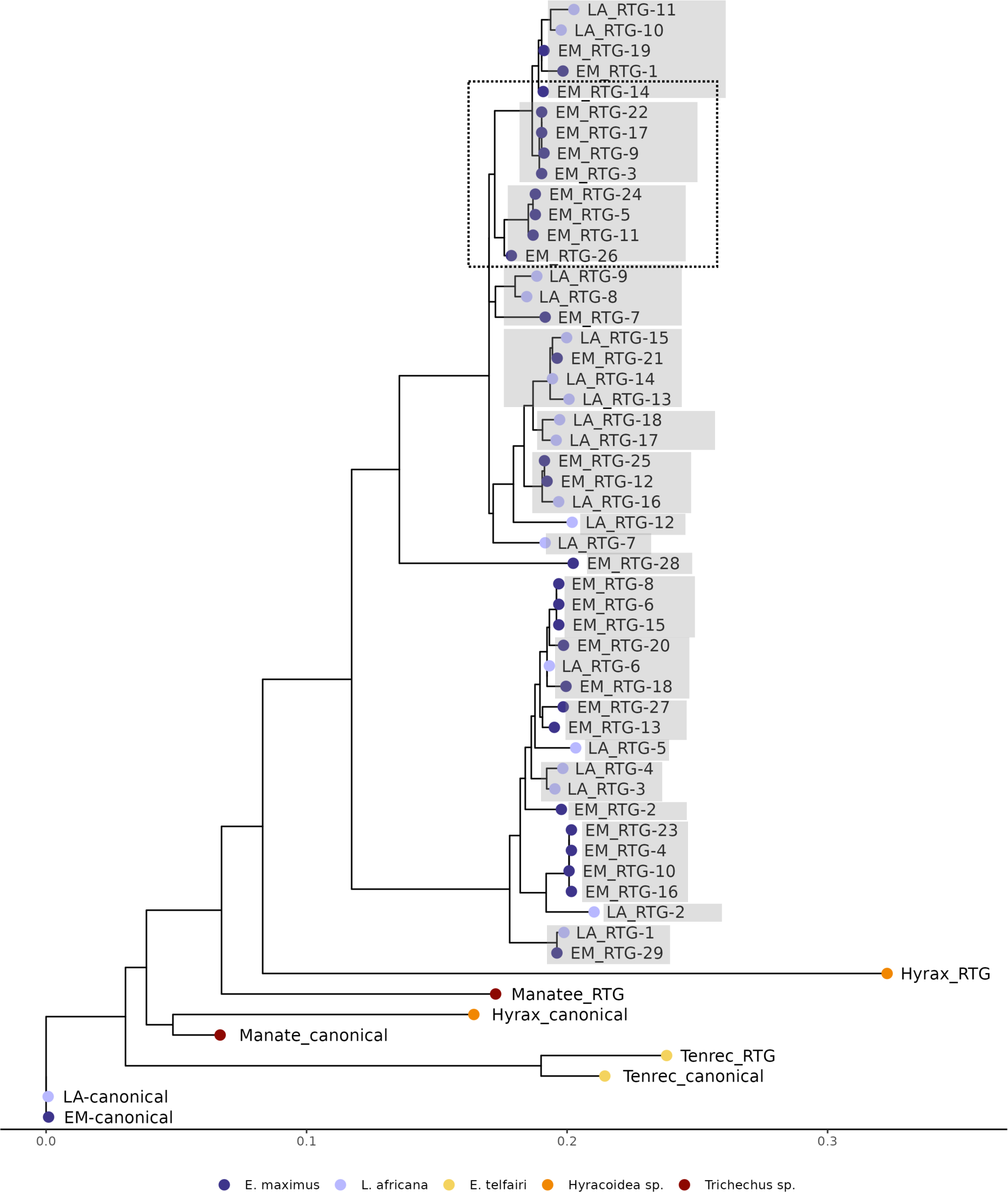
Phylogenetic analysis of the *TP53* RTGs in African and Asian elephants and phylogenetically related species. EM: *E. maximus*; LA: *L. africana*). Each coloured dot represents the species to which each RTG belongs to, as specified in the legend. The multiple copies in the dotted rectangle suggests concerted evolution between the copies of each species and multiple gene conversion events.

## Notes

### Competing Interest Statement

The authors have declared no competing interest.

